# Identifying the genetic and epigenetic basis for asymmetric bZIP expression in temperature-stressed bread wheat

**DOI:** 10.1101/2024.01.04.574129

**Authors:** Raminder Kaur, Dalwinder Singh, Pankaj Kumar, Gazaldeep Kaur, Koushik Shah, Harshita Pandey, Shamjetsabam Gangarani Devi, Ajay Kumar Pandey, Vikas Rishi

## Abstract

Asymmetric expression in the bread wheat (*Triticum aestivum*) genome refers to the differential expression of genes from A, B, and D parental genomes. Bread wheat is a hexaploid crop with six copies of each chromosome. This complexity can result in unequal expression of genes from each parental genome, leading to asymmetry in gene expression. In other polyploid crops like cotton, transcription factors (TF) exhibit genome-biased expression; however, there are no comparable studies for bread wheat. One of the most prominent TFs families in plants is the basic Leucine Zippers (bZIP) which are eukaryote-specific proteins and regulate various biological processes, including stress-related responses. bZIP proteins are dimeric and several heptads long. They exhibit typical coiled-coil structures with strategically placed amino acids in each heptad, responsible for their stability and specificity. Here, we aim to decipher the structural basis of the asymmetric expression of the bZIP TFs in wheat under low and high-temperature conditions. Furthermore, 19 highly expressed stress-related TabZIP TFs were analysed for their asymmetric expression profiles as plants were exposed to temperature-stress conditions. Two benchmarks were used to analyse the asymmetric gene expression of bZIPs, i.e., a) the promoter’s occupancy by the epigenetic marker histones, namely, H3K4me3, H3k9ac (active) and H3K27me3 (repressive), b) density and diversity of cis-regulatory elements in the promoters. Notably, the genetic basis of the differences in protein sequences of bZIP triads was explored, which may impart structural stability to a specific homeolog enabling the plant to endure the stress conditions better.

## Introduction

The expression of genes from each parental genome in a polyploid plant differs and is known as asymmetric gene expression. Based on subgenome expression, alleles may be balanced, dominant, or suppressive. Like other polyploid crops, sugarcane and cotton, Triticum aestivum, a hexaploid crop, exhibits genomic biases. It contains 42 chromosomes with contributions from three, i.e., A, B, and D genomes. One of the main challenges in studying asymmetric gene expression in wheat is distinguishing between the expression of homoeologous genes, which are nearly identical genes located on different chromosomes (Ramírez-González *et al*. 2018). This is particularly important in wheat, as the hexaploid genome has many homoeologous genes, making it challenging to identify which gene is expressed from which parental genome. Nevertheless, recent advances in RNA sequencing technology and computational tools have made it possible to accurately quantify the expression of homoeologous genes, allowing researchers to study asymmetric gene expression in wheat and other polyploid crops (Kaur *et al*. 2023). For example, RNA sequencing using "digital gene expression" (DGE) has been used to accurately quantify gene expression from each parent genome in the hexaploid wheat (Xiao *et al*. 2013). In addition, computational tools have been developed to identify and distinguish between homoeologous transcripts, allowing for the accurate quantification of asymmetric gene expression (Frazer *et al*. 2004). The mechanistic basis of allele-specific differential gene expression is explained based on the active H3K4me3 and H3K9ac and suppressive H3K27me3 histone modifications and the DNA methylation status of CG, CHH, and CHG bases in the promoter region, and the gene body (Ramírez-González *et al*. 2018; Xue *et al*. 2022). Additionally, the activity of an allele is determined by variations in its core cis-elements in the promoter region (Zhao *et al*. 2018). In a classical study, previous research with cotton showed that the asymmetric gene expression of *PRE1*, which codes for a Basic Helix Loop Helix (bHLH) TF, leads to fiber lengthening (Zhao *et al*. 2018). To our knowledge, no similar TF-associated study involving allele-specific expression on wheat has been undertaken.

Pathogens, herbivores, cold, salt, drought, flood, extreme temperatures, and toxic heavy metal ions are a few of the many biotic and abiotic stresses that plants routinely encounter due to their sessile nature (Alves et al. 2013, Schütze et al. 2008). Activating stress-responsive genes is one of the adaptive strategies plants have developed to deal with these stressful environmental situations (Latchman 1997; Fujita *et al*. 2006; Cheng *et al*. 2011). bZIP (basic leucine zipper) transcription factors (TFs) are the critical regulators of this process, acting as sensors of stress signals and mediating the expression of stress-responsive genes. bZIP plays a crucial role in plants’ stress response. For example, ABF2 in *Arabidopsis* and OsbZIP23, OsABI5, OsbZIP46, TRAB1 in rice, ZmbZIP23, and ZmbZIP112 in maize are known bZIPs involved in ABA signalling and regulation of plant fertility and drought tolerance (Jakoby *et al*. 2002; Nijhawan *et al*. 2008; Ying *et al*. 2012). ABI5 is involved in the ABA signalling pathway, plant growth metabolism, and various stress responses (Lin *et al*. 2021). VIP1, an *Arabidopsis* protein, known to interact with Agrobacterium VirE2 protein, and is involved in VirE2 nuclear import and Agrobacterium infection activity (Tzfira *et al*. 2001).

Hexaploid bread wheat with a 17Gb genome size is predicted to code for 294 bZIPs, (Appels *et al*. 2018), some of which are known to be involved in mitigating heat, drought, and temperature stress. *TabZIP6* reduced the freezing tolerance of transgenic *Arabidopsis* seedlings by downregulating the expression of *CBFs* (Cai *et al*. 2018). *TaWLIP19* bZIP is upregulated in wheat under salinity conditions (Baloglu *et al*. 2014). An abiotic stress structure-function link has been postulated using a molecular model of wheat bZIP bound to a DNA (Sornaraj *et al*. 2016). Hexaploid or bread wheat has three homeolog of each bZIP TFs that show subtle changes in amino acid sequences. All alleles are anticipated to contribute to the expression of a specific bZIP. This is challenged by the new finding demonstrating allele-specific gene expression (Ramírez-González *et al*. 2018; Zhao *et al*. 2018), but the concept of such asymmetric gene expression is poorly understood. Using *in silco* methodology and expression analysis, we have shortlisted 19 bZIP TFs, which are misregulated under temperature stress conditions.

Structurally, The bZIP protein contains an N-terminal basic amino acid-rich DNA-binding domain followed by a leucine zipper region containing amphipathic peptide with a high propensity to form a coiled-coil dimer (Vinson *et al*. 1989, 2006). Embedded in the leucine zipper regions are short sequences called ‘Trigger sequences’ that facilitate helix formation in bZIP monomer (Steinmetz *et al*. 2007). The trigger sequence’s ability to induce dimer formation in the bZIP protein may alter subtly with the change in single amino acid. Here, we asked if expressing a specific allele gives a distinct advantage to plants when faced with environmental stress conditions. This will provide a novel structural insight into genome-biased expression.

## Materials and methods

### Identification of the stress-related bZIP genes in bread wheat and chromosomal location analysis

The hidden Markov model of the bZIP domain (PF00170) was downloaded from the Pfam (protein domain family) database, and the bZIP domain’s presence was inspected in the stress-responsive TabZIPs. After reviewing the abiotic stress-related bZIPs from available literature, 19 stress-related *TabZIP* were identified for subsequent study. The information on the plant species’ CDS and protein sequences of *T aestivum*, *H vulgare*, *O sativa*, *Z mays,* and *A thaliana* were used from the *EnsemblPlants* database (Supplementary Table 1). The selection of putative abiotic heat and cold stress-related genes was confirmed by conceiving the gene expression information from the wheat expression browser named expVIP tool (Supplementary Table 2). The *TabZIPs* were named with the transcript IDs (Traes IDs) as given in the *EnsemblPlants,* representing a unique identifier for each bZIP transcription factor. Along with a few earlier reported bZIP files, the listed names and transcript IDs are also included. In addition, the coding sequence length, amino acids composition, location on the chromosome, uniport ID, e-value, percentage ID, and reference genome or assembly information were retrieved for each bZIP and are listed in the Supplementary Table1. Domain analysis revealed that all but five potential bZIP transcription factors had a typical bZIP domain with an invariant N-_x7_-R/K (where x is any amino acid) motif in the basic region and a Leu or other bulky hydrophobic amino acids at the ‘d’ position of the heptad nine amino acids c-terminus downstream of R/K. To a large extent, the Leu or other hydrophobic amino acid pattern is repeated in each n heptad length (3-14). The leucine zipper sequences contain the typical N conserved at the ‘a’ position, and its signature pattern largely is responsible for the dimerization behavior of a bZIP. Further, the MapChart tool was used to map the chromosome location of these stress-related *TabZIPs* (Supplementary Figure 1).

### *TabZIP* structure and promoter enrichment analysis in hexaploid wheat

CDS sequences of *TabZIP*s were submitted for gene structure analysis. The gff3 files were used to extract the intron-exon, UTR (non-coding region) distribution, and other gene structure information. They were finally visualized in GSDS server (Hu *et al*. 2015). MEME software was used to analyze the conserved motifs in the proteins (Bailey *et al*. 2015). The maximum number of motifs found was 10, and motif width was 6–20, as the other parameters were set as default values. Promoters affect gene function to a large extent. The promoter region contains potential transcription factor binding sites which superintend transcription factor-mediated gene regulation. The 2000 bp upstream sequences from the transcription start site (TSS) were submitted to the PlantCARE tool to analyze the bZIP promoters for cis-element enrichment (http://bioinformatics.psb.ugent.be/webtools/plantcare/html) (Supplementary Table 3).

### Duplication, Ka/Ks, and Synteny Analysis

To check the evolutionary-derived effects on the gene expression, collinearity analysis was performed for the complete TabZIP family members (294) to visualize the collinearity between wheat and reference plant species’ syntenic and non-syntenic gene pairs. The Blastall tool was used with the e-value=1 × 10^−20^ (Mount 2007). MCScanX tool was used to analyse the gene duplication and synteny events. The CDS sequences were used to bring forth the evolutionary relationship between bZIP genes by synonymous substitution rate (Ks) and nonsynonymous substitution rate (Ka). Ka/Ks ratio in wheat was calculated using KaKs calculator (Yang and Nielsen 2000) (Supplementary Table 4). Duplicated gene pairs were classified into four types: tandem, proximal, segmental, and dispersed duplications. Circos tool was used to visualize the collinear regions heightened with the names of TabZIP members. The divergence time of gene duplication (MYA) was calculated by the formula T = Ks/2λ with a value of =6.5 × 10^−9^ synonymous substitutions per site per year (Lynch and Conery 2000).

### Expression analysis based on RNA-seq

The expression levels for the candidate bZIPs were extracted across developmental stages and tissue types, in addition to the changes in their expression in abiotic stress conditions. Normalized gene expression values across numerous development stages were obtained from expVIP (http://www.wheatexpression.com/). In addition, log fold change (LFC) values were calculated for the stressed conditions w.r.t. controls. Gene expression log_2_ values >1 were treated as upregulated, whereas those with <1 were considered downregulated. The expression levels and LFC values were used to plot heatmaps using the Heatmap2 package in R.

### Plant material and treatment

The hexaploid bread wheat variety ‘Chinese spring’ was used for all experiments. For temperature stress experiments, ten-day-old seedlings were given stress in growth chambers set at required temperatures, i.e., 4 °C, 37 °C, and 42 °C for 48 hrs. Controls were kept at ambient conditions (24 °C). After treatment, seedlings were harvested for RNA isolation and kept at −80 °C till further processing. Total RNA was extracted from the leaves of unstressed and stressed wheat seedling samples at 0, 1, 6, 24, and 48 hrs.

### Total RNA isolation and qRT PCR

Samples’ total RNA was manually isolated using TRIzol^®^ Reagent (Invitrogen, Canada). Subsequently, DNase treatment was given to purify the RNA samples from DNA. Quality assessment was conducted using a NanoDrop ONE spectrophotometer (Thermo Fisher Scientific, USA). RNA samples were checked for any degradation on 1.2% agarose gels. 2.0 µg of total RNA from each sample was used for cDNA preparation. iScript™ cDNA Synthesis Kit (Bio-Rad, USA) was used to prepare cDNA samples per the manufacturer’s protocol. The primers were designed in the PrimerQuest™ tool (Integrated DNA Technologies, USA) by following default parameters.

The real-time PCR reactions were carried out in C1000 Touch CFX 96 real-time Thermocycler (Bio-Rad, USA) for 35 cycles. Three biological replicates were considered for each sample. The data were normalized using *actin* as an internal control. The relative expression was estimated using the 2^-ΔΔCT^ method. The primers used are listed in Supplementary Table 5.

### Statistical Analysis

GraphPad Prism 7.0 statistical software was used for data analysis, and Student’s t-test was employed to test significance, * *p* < 0.05 (Supplementary Figure 2). All experiments were repeated 3-5 times. The values are the mean ± one standard deviation (SD) of three biological replicates. Data presented in this study represent three biological replicates with 5–10 seedlings per experiment.

### Analysis of wheat bZIP homeologs divergent expression pattern

Homeolog Triads taken in the study are closely related gene copies of three parent A, B, and D sub-genomes. Transcript expression values and physical coordinates were fetched from the expVIP browser. The expression profiles of 19 selected bZIPs were evaluated, considering the presence of active or repressive epigenetic markers in their promoter regions. We spotted the divergent expression trend in the ChIP-seq data available from Gene Expression Omnibus (*GEO*) under accession no. GSE139019. Triads representing the altered genome expression trend were used, and two genes out of 19 showed two instead of three homeologs. As mentioned, ChIP-seq data were used for specific temperature stress conditions where biological duplicates were exposed to heat and cold stress.

Further, we have performed genome-wide mapping of hexaploid wheat genes on each chromosome. Such an analysis led to the physical location of candidate genes in the distal and proximal regions of chromosomes to further catechize the dynamic gene expression. Next, we have considered the amino acid sequence differences within the bZIP homeologs, giving prominence to each heptad’s ‘abcdefg’ amino acid composition. The amino acid differences among the homeologs potentiate their structural stability and dimerization potential important for their function under stress conditions.

## Results

### Identification of the *bZIP* family in wheat

In all, there are 294 TabZIPs listed on *EnsemblPlants*. After reviewing the abiotic stress-related bZIPs from available literature, 19 stress-related *TabZIP* were identified for subsequent study. The nomenclature of these genes was given based on the Traes IDs available in *EnsemblPlants*, which includes the parent subgenome information. The primary physiological and physicochemical data of the candidate *TabZIP* was analyzed, including the protein sequence lengths, relative molecular weight (MW), and various physicochemical properties. As shown in Supplementary Table 1, the amino acids length of the coding proteins ranges from 150 (TraesCS1A02G072600) to 649 amino acid residues (TraesCS1B02G268500), and the average protein length is 339 amino acids. The isoelectric point of the proteins ranged from 4.7 (TraesCS7B02G299200) to 10 (TraesCS7D02G349300). The molecular is in the range of 16284 (TraesCS1A02G072600) - 68700 Da (TraesCS1B02G268500), with an average value of 33,878 Da (Supplementary Table 1). The subcellular location showed all *TabZIPs* to be in the nucleus, reflecting the gene family’s biological function as transcription factors. http://bioinfadmin.cs.ucl.ac.uk/index.html

### Gene structure and motif analysis of *TabZIP*

We obtained the exon-intron arrangement in the identified *TabZIP* by screening the coding and genomic DNA sequences. To better understand the *TabZIP* family in bread wheat, structural analysis was performed by comparing each *bZIP* in wheat to understand structural diversity. The results showed that 15 of 19 temperature stress-related *TabZIP* have introns, which were closely related (Figure 1a). All stress-related *TabZIP* contained exon-rich coding regions. Regular motif distribution was observed with 1-8 motifs (input settings). There are 4 bZIPs (TraesCS5A02G237200, TraesCS3D02G364900 (ABI5), TraesCS7D02G171300, and TraesCS6A02G333600) that show additional motifs. Using the MEME program, 12 of 19 *bZIPs* have exclusive bZIP motifs present, and included TraesCS5D02G124600 (TabZIP1) (Figure 1b).

**Figure 1:**
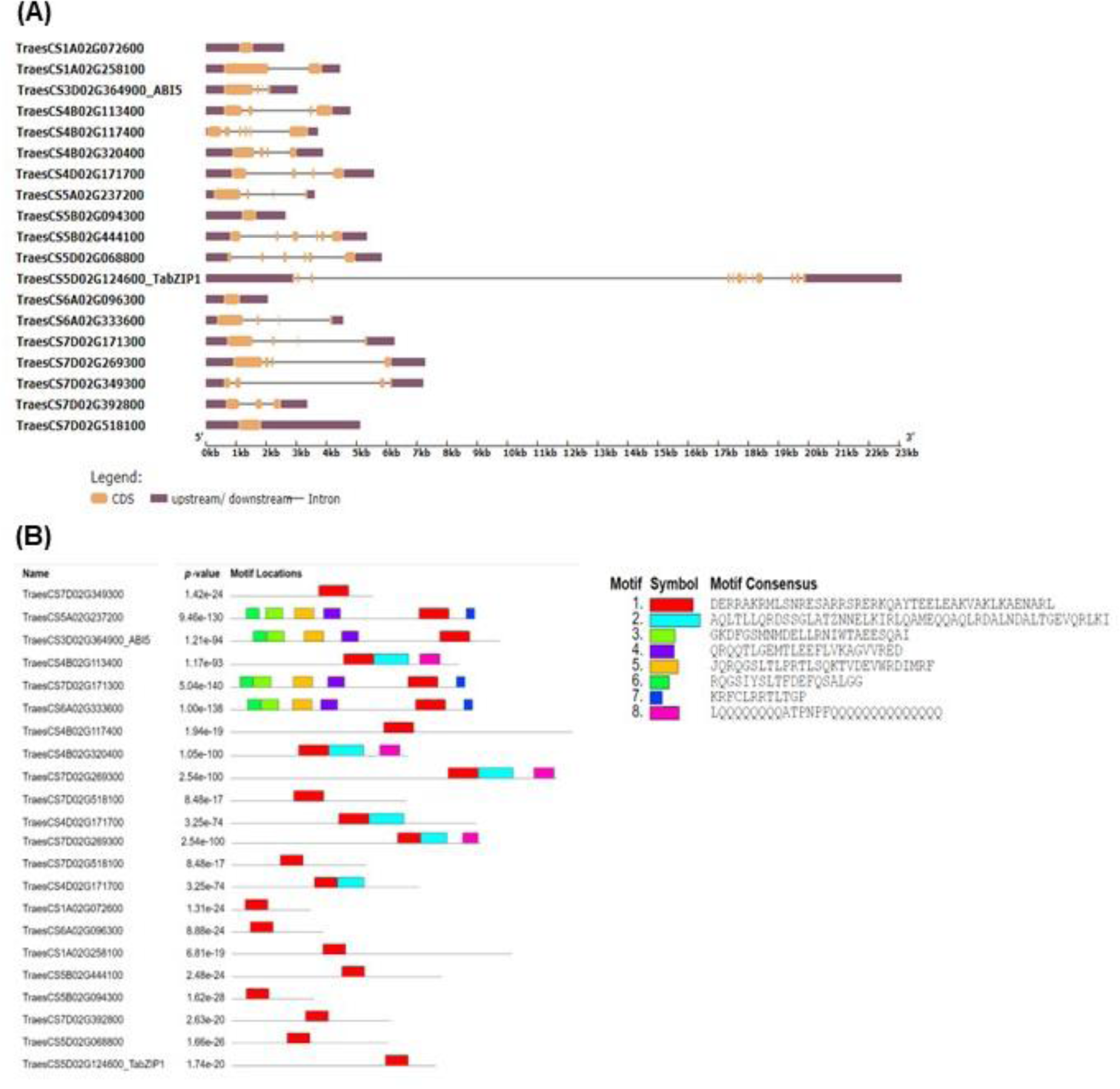
Distribution of gene structure and conserved motifs present in the candidate genes. (**A**) Representation of gene structure of the *T.aestivum* bZIP gene family in the candidate genes with yellow, purple boxes and black horizontal lines displayed the Coding sequence (CDS), Untranslated regions and introns respectively. (**B**) Location of Conserved motifs located on each gene displayed with the color box details on the right side.

### Chromosomal localization and synteny analysis of *TabZIP* genes

All the stress-related *TabZIPs* were found on 18 of 21 chromosomes covering around 91% of the whole genome (Supplementary Figure 1). Some bZIP genes are present on each chromosome, although their distribution varies greatly and is uneven. For example, chromosomes 5 and 7 have the most *TabZIP* (∼53%), which is absent on chromosome 2. Chromosome 4 contained four members (21%). Further, chromosomes 1 and 3 have 2 and 1 *TabZIP*, with a 10% stress-related share in the genome. Moreover, most of the genes were distributed near the ends of the chromosomes. Besides, to understand the evolutionary relationship of the *bZIP* family in wheat, collinearity relationships were displayed by comparing wheat with four other species. These species include a dicotyledon (*Arabidopsis*) and three monocotyledons (barley, maize, and rice) (Figure 2a).

**Figure 2:**
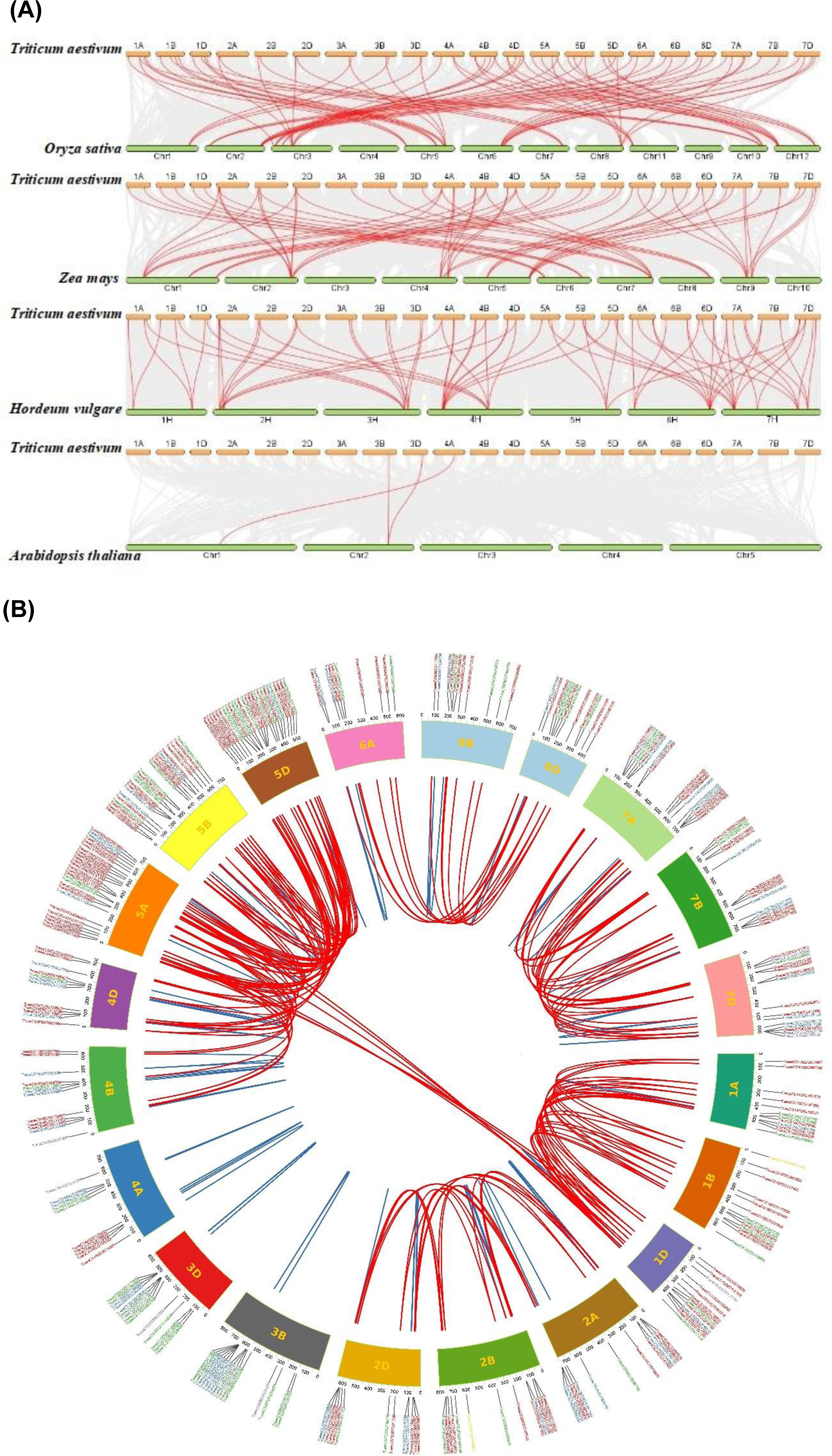
Synteny analysis of *T aestivum* with reference plant species and syntenic pairs of *TabZIP* family following different modes of gene duplication (**A**) Synteny analysis of *TabZIP* genes between *Triticum aestivum* and four representative plant species such as Oryza sativa, Zea mays, Hordeum vulgare and Arabidopsis thaliana. Gray lines indicate the collinear bars within the *Triticum aestivum* and other plant genomes, whereas the red lines highlight the syntenic *TabZIP* gene pairs. (**B**) The syntenic pairs of wheat *TabZIP* genes from different duplication events. Corresponding color lines linked syntenic pairs from different duplication modes. Red, blue, purple, and green represent syntenic regions with segmental, tandem, transposed, and proximal duplication. Other syntenic pairs genes were shown with pale yellow lines.

### Expansion and evolutionary analysis of *TabZIP* family

The duplicated gene pairs in this study were all identified based on the duplication events in *T. aestivum*, such as Tandem Duplication (TD), Proximal Duplication (PD), and Whole Genome Duplication (WGD). About 53% of the duplicated genes were identified as WGDs (165 gene pairs), and 22 % (70 gene pairs) were identified as Proximal Duplicate gene pairs. Additionally, there is a 20 % occurrence (63 gene pairs) of Tandem Duplicated genes, and the remaining dispersed and singleton gene pairs accounted for 4 % and 1 %, respectively (11 and 2 gene pairs, respectively). Only two gene pairs show a singleton type of duplication event, meaning these genes may have strongly adapted during the selection pressure (Pace *et al*. 2009). The non-synonymous mutation rate to the synonymous mutation rate (Ka/Ks) is essential to determine the selection pressure. Therefore, the YN method was followed to calculate the Ka and Ks values. The results showed that the Ka/Ks ratios of stress-related TabZIPs were <1, explaining the purifying selection during evolution (Supplementary Table 4 and Figure 2b). It is predicted that allohexaploid bread wheat *T. aestivum* (AABBDD) was formed 8500-9000 years ago by a hybridization event between tetraploid progenitor (AABB) and diploid donor (DD). To better understand the origin of stress-related *TabZIPs* and gain insight into transcriptomic changes due to genome polyploidy, we conducted the collinearity analysis between *T aestivum* and other species. As a result, we obtained the orthologous genes in *A thaliana*, *O sativa*, *Z mays*, and *H vulgare*. The Ka/Ks ratio is less than 1, confirming that *T aestivum* has undergone a stringent purifying selection.

### Cis-acting element analysis

Transcription factors (TFs) liaise with cellular responses mediated by binding to specific cis-acting DNA sequences in the promoter regions. As sessile organisms, plants develop and encounter various environmental stresses. Hence, signalling cascades govern the development, and the stress switches converge at the gene expression level. Binding transcription factors at cis-elements in plant promoters is a decisive step in gene expression and regulation. As a result, considerable effort has been expended in dissecting various plant promoters, identifying multiple transcription factors and specific interactions with other TFs, and exploring their functional aspects. Promoter cis-element analysis makes it easier to comprehend gene expression and function. Many cis-elements are in the upstream sequence of stress-related *TabZIP* (Figure 3). In addition, cis-elements, such as TATA-box, TGACG-box, and CAT-box, are related to transcription initiation, growth, and development. Many photo-responsive elements, such as AS1, W-box, A-box, G-box, and endoplasmic reticulum (ER) stress-related TGACG motif, suggest these TFs function in the development. Furthermore, CGTCA-motif (methyl jasmonate-induced), ABRE (ABA response element), and as-1 (Auxin response element) are responsive to environmental cues. However, there are common cis-elements associated with abiotic stresses response, e.g., ABRE (abscisic acid-responsive element), low-temperature response (LTR), involved in dehydration stress (DRE), drought (MYB, and MYC), osmotic stress response anaerobic inducible enhancer (GC-Motif), and antioxidant response (ARE). Ultimately, consideration was given to the cis-acting regulatory sites in the promoter of *TabZIP* that are abundant in stress-related sequences, especially those related to heat and cold temperature stress.

**Figure 3:**
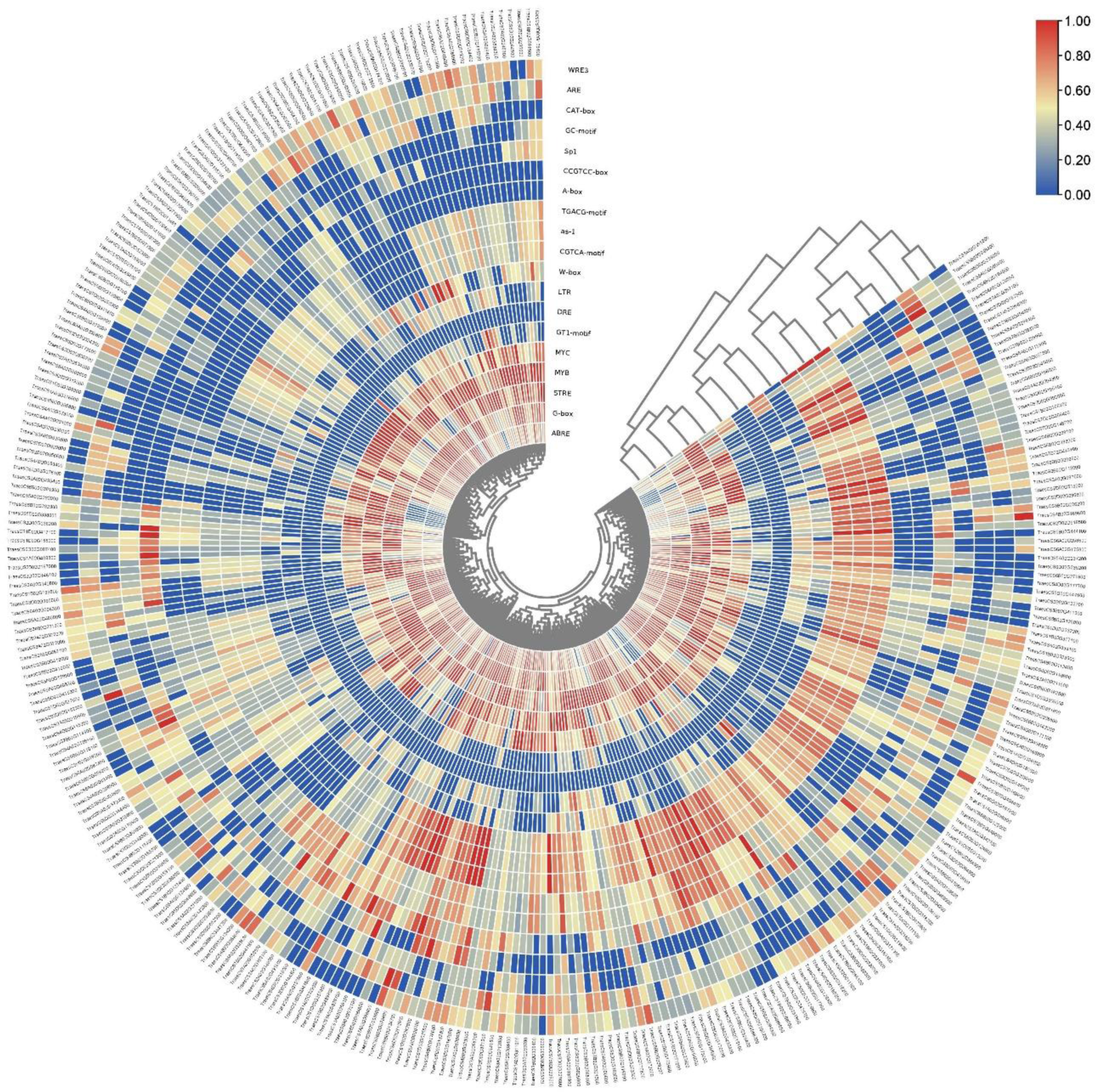
Predicted cis-elements in the promoter region of *TabZIP* family. The analysis was carried out by the Plantcare tool with 2000 bp upstream to the transcription start site (TSS). Putative cis-elements were highlighted by different colors and labelled. The depth of the color represents the number of times the cis-element occurs, and a detailed color scale is included.

### Expression study of *TabZIPs* associated with stress under diverse conditions and developmental phases

The expression profiles from RNA-seq data from expVIP were explored under various abiotic stress conditions such as drought, heat stress (DsHs), and cold stress. In addition, transcriptional levels of *TabZIP* in grain tissue development (GTDT), development time course of Chinese spring cultivar (DTCS) across other tissues, and in reproductive organs development such as stamen, pistil and pistillody (SPP) (Figure 4). Log fold change values were used to plot heat maps for all developmental stages. Among them, red cells represent the up/high-regulated gene expression with log2 fold change (LFC) values >1 (significantly upregulated) and blue ones with LFC<1 (significantly down-regulated) across the tissue development.

**Figure 4:**
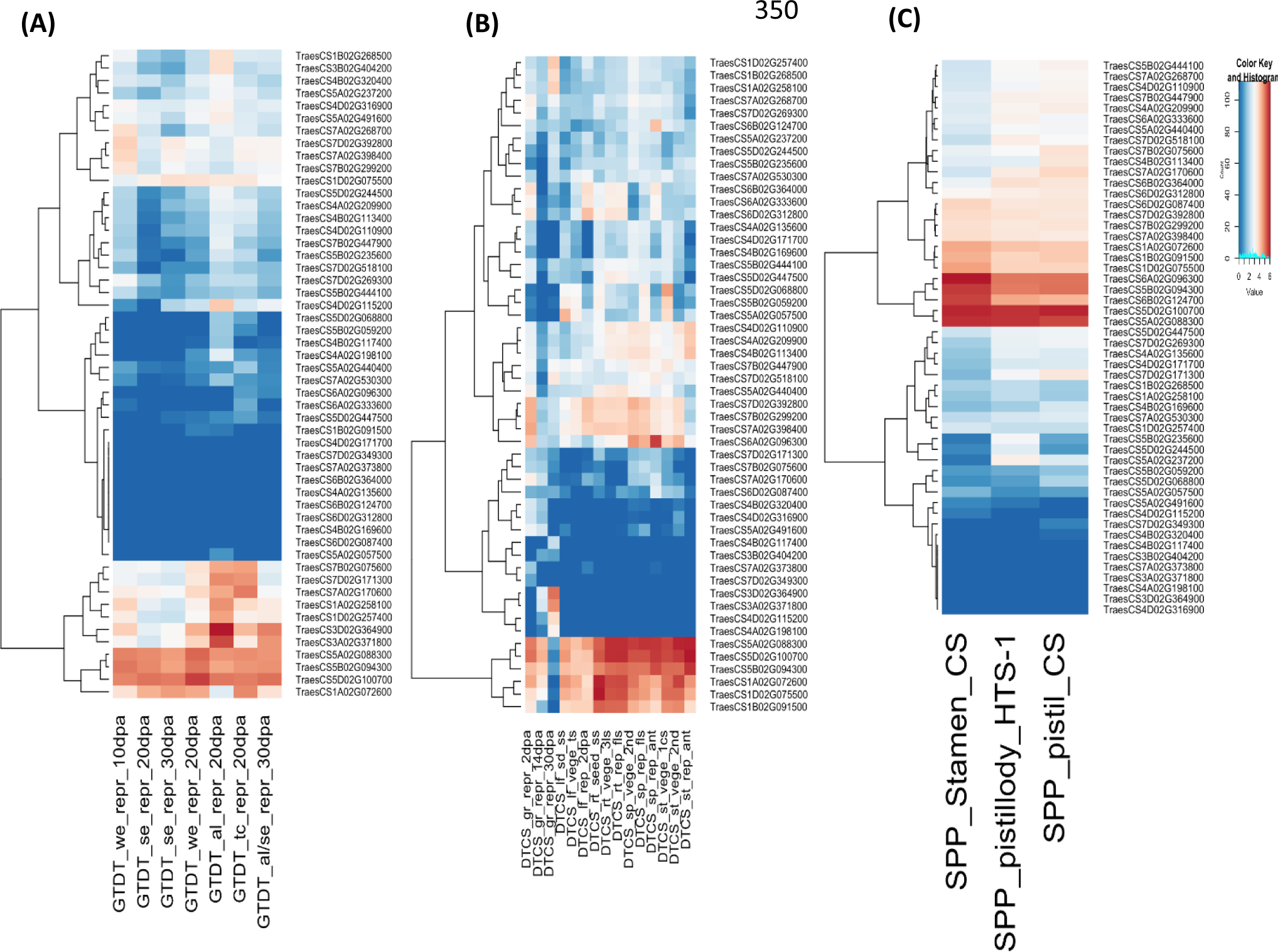
The expression levels for the candidate *TabZIP* genes were extracted across different developmental stages and tissue types in *Chinese spring* cultivar. (**A)** This section contains expression level of candidate genes with homeologs with respect to (w.r.t) to normal grain tissue developmental time course. (**B)** This section shows the expression level of developmental time course of Chinese spring cultivar. (**C)** It specifically represents the expression level of candidate genes with their homeologs in the reproductive tissue types such as stamen, pistil and pistilloidy. TPM counts were [http://www.wheatexpression.com/]) for expression across numerous stages of development. The TPM values were used to plot heatmaps using heatmap2 package in R. Detailed Color scale has been shown on the right side of the image.

qRT-PCR validated the expression study related to temperature stress in seedlings. 19 *TabZIP* genes were selected based on the publicly available expVIP database expression database for qRT-PCR validation. In ExpVIP, several wheat varieties show the expression details with specific cultivars possessing the highest transcript levels while encountering stress conditions. Information on gene-specific primers used in the study is given in Supplementary Table 5. Gene expression analyses were performed in Chinese spring cultivars, focusing on the 10-day-old seedling stage of plants. The expression profiles of 19 *TabZIPs* under abiotic stress conditions were examined in 10-day-old seedlings subjected to temperature stress at 4 °C, 37 °C, and 42 °C for 1, 6, 24, and 48 hrs.

TraesCS5B02G094300 (along with its homeologs) has shown the highest expression of 7.65 log_2_FC (LFC) in the stamen, pistil, and pistillody stages of development. In the seedling stage, TraesCS6A02G096300 and TraesCS1A02G072600 (and their respective homeologs) exhibit a similar expression trend in the above stages as demonstrated in an earlier study (Chauhan *et al*. 2011). TraesCS4B02G320400 showed an enriched expression of 6.80 LFC in heat stress at 42°C in 1 hr of exposure. In the DTCS and GTDT development stages of seeds, *TabZIPs* such as TraesCS7D02G171300, TraesCS3D02G364900, TraesCS5B02G094300, and TraesCS1A02G072600 are present with high transcript level. TraesCS5B02G444100 shows an increased expression of 9.13 LFC till 24 hrs of 37°C exposure. After 48 hrs of heat exposure, TraesCS6A02G096300 has a high expression level of 9.41 LFC at 42°C, and in addition, its fold enrichment is 5.45 under freezing stress. TraesCS3D02G364900 (ABI5) is an important TabZIP, which is downregulated explicitly in heat stress with -5.92 LFC at 37°C (after 1 hr), - 2.300 at 42°C (after 6 hrs) but increased to 9.23 LFC at 4°C even after 24 hrs of freezing stress. Furthermore, TraesCS1A02G258100 shows an average expression in grain tissue development. More compelling grounds exist to investigate when these putative TabZIPs are populated in the grain (the main application is wheat) and other high-expression tissue development stages. The expression of the 10 of 19 genes (TraesCS3D02G364900 (ABI5), TraesCS4B02G113400, TraesCS7D02G171300, TraesCS5A02G237200, TraesCS4B02G117400, TraesCS7D02G269300, TraesCS7D02G518100, TraesCS1A02G258100, TraesCS5D02G068800 and TraesCS5D02G124600 (TabZIP1) ) were downregulated in the heat stress condition at 37°C. Thereafter, only TraesCS5A02G237200 and TraesCS5D02G068800 remained significantly downregulated at 42°C. Rest 8 are induced by heat stress (37°C and 42°C). Besides, TraesCS4B02G117400 shows a -7.674 LFC in the freezing stress, i.e., exposed to 4°C (Figure 5). In qRT-PCR experiments, we failed to observe the expression of TraesCS5A02G491600, TraesCS4D02G171700, and TraesCS7D02G392800. After qRT PCR validation, we considered investigating the biased or asymmetric expression pattern in the *TabZIP* homeolog copies. Homeolog-specific primers were designed (Supplementary Table 5) to further investigate the asymmetric expression of a bZIP (TraesCS1A02G072600, TraesCS1B02G091500, and TraesCS1D02G075500). Supplementary Figure 5 shows the expression patterns of the A, B, and D homeologs under three different temperature stress conditions (4°C, 37°C, and 42°C). There is evidence of a homeologs-specific expression pattern. Homeolog B and D are highly expressed at 37°C and 42°C, respectively, while homeolog A is mostly expressed at 4°C. Homeolog-specific primers were designed (Supplementary Table 5) to further investigate the asymmetric expression of a bZIP (TraesCS1A02G072600, TraesCS1B02G091500, and TraesCS1D02G075500) in the seedling stage. Supplementary Figure 5 shows the expression patterns of the A, B, and D homeologs under three different temperature stress conditions (4°C, 37°C, and 42°C). There is evidence of a homeologs-specific expression pattern. Homeolog B and D are highly expressed at 37°C and 42°C, respectively, while homeolog A is mostly expressed at 4°C (Supplementary Figure 5).

**Figure 5:**
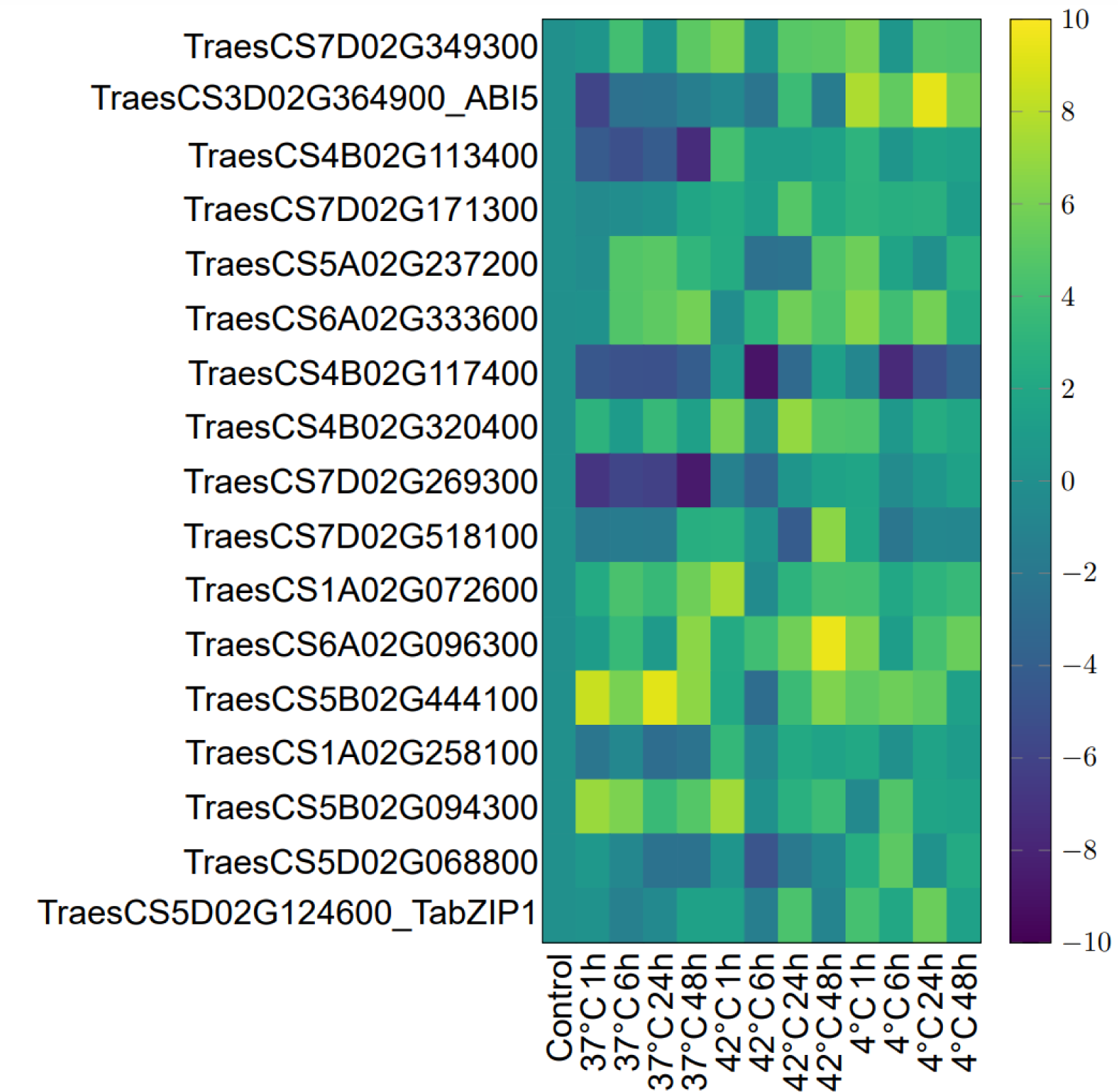
Heatmap representing quantitative real-time PCR validation of 19 candidate *TabZIPs* in 10 day old seedlings exposed to three temperature (cold and heat stress) conditions (4 °C, 37 °C and 42 °C) for 1,6,24, and 48 hours. 24 °C (physiological room temperature) was taken as control temperature. The color scale indicates the fold-change value converted to log2 scale, as compared with untreated sample. Values are expressed as mean ± SD; *n* = 3 independent experiments, and transcript was normalized to β-actin as internal control gene.

### The genetic and epigenetic basis of the temperature stress-induced asymmetric expression of wheat *bZIP* homeologs

Large and complicated genomes, like those of cotton (Yoo et al. 2013) and bread wheat (Appels et al. 2018), make it difficult to examine the gene regulation of polyploid agricultural plants. We herein followed the genetic (sequence-based) architecture and the presence of epigenetic markers in the upstream promoter regions for exploring the prejudiced expression between the homeologs, followed by analysing structure-based advantage a specific homeolog may confers to the plant under stress conditions.

We have mapped the entire bread wheat genome with gene expression TPM values on seven chromosomes and highlighted the 19 candidate genes with 55 homeologs in circos plots (Figure 6). All the 19 candidate genes and their homeologs mapped to the distal regions in the respective chromosomes, suggesting a dynamic and altered gene expression (Figure 6). The asymmetric expression pattern has been observed in the candidate homeologs of bread wheat. The study is the first attempt to decipher the structure-function basis of the condition-specific expression of a homoeolog. In a bZIP, the leucine zipper region contains several heptads repeats having charged, hydrophobic, and polar amino acids at ‘abcdefg’ positions. Bulky amino acids in ‘a’ and ‘d’ positions are buried and provide stability to the structure, whereas amino acids at ‘g’ and ‘e’ positions are often charged and contribute specificity to dimer formation. ‘b’, ‘c’, ‘f’ positions have polar amino acids and are exposed. The individual amino acid at ‘a’ and ‘d’ contribution to stability is ranked by coupling energy (Moitra, Acharya). In addition, amino acids at ‘b’, ‘c’, ‘f’ positions in a heptad may help bZIP interact with other proteins (Lin Chen, Nature, 1998). It is proposed that monomers that lack structure come together during dimer formation to form a coiled coil. It is also reported that in monomers, short sequences called “Trigger sequences” nucleate and greatly facilitate alpha helices formation, thus initiating a coiled coil (Steinmetz *et al*. 2007). Of 19 bZIP proteins, 10 showed no difference in expression among homeologs, whereas 9 showed homolog-specific expression. Of 10, 7 bZIPs homeologs showed no change in amino acid sequences, whereas 3 have different sequences. 9 bZIPs with genome-biases have 5 proteins whose amino acid sequences differ. 4 bZIPs homeologs have the same protein sequence (Figure7 and Supplementary Figure 4). In this context, we found that TraesCS5B02G235600 homeolog differs from the other two homeologs by having negatively charged ‘E’ at the ‘b’ position in the second heptad and positively charged lysine at ‘b’ position in the third heptad. Such substitution led to several strong interactions in this homeolog compared to the other two, suggesting a trigger-like sequence. Glutamic acid and lysine at ‘b’ positions induce stronger interactions with other residues, which may lead to a more stable alpha helix structure in monomeric conformation (Supplementary Figure 3). Also, the leucine at ‘a’ position in TraesCS3A02G371800 in the fifth heptad has a lower coupling energy than asparagine at the same position in other homeologs, therefore, is a more stable (Acharya *et al*. 2006). Similar observations are made for TraesCS3B02G404200.

**Figure 6:**
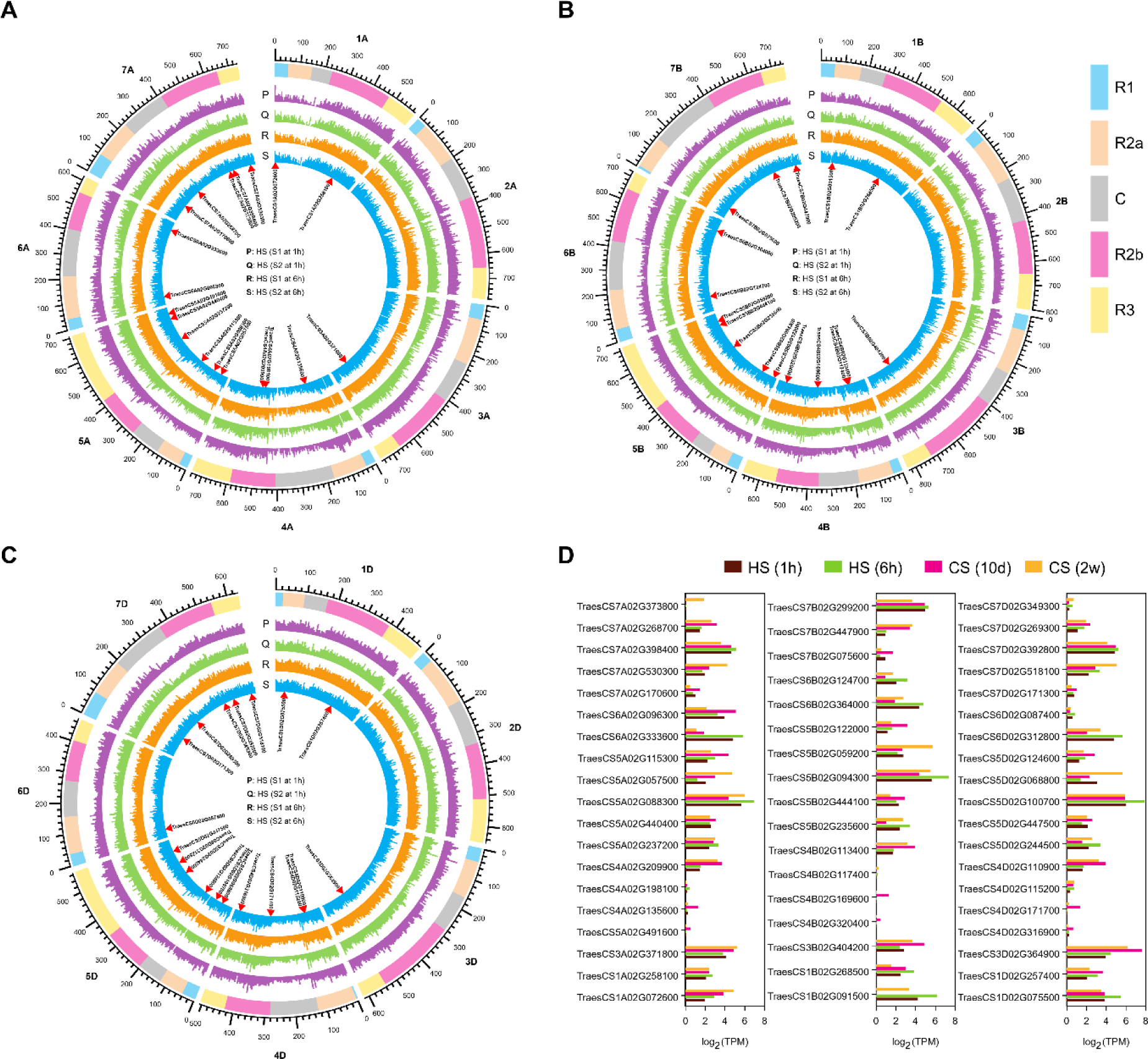
Genome-wide expression of all genes in T. aestivum on the seven chromosomes of each subgenome (**A**) shows genome wide expression of T. aestivum protein coding genes in the A-subgenome for two heat stress replicates at 1h (P,Q) and 6 h (R,S) which were obtained from study (https://www.ncbi.nlm.nih.gov/sra/SRP045409). Similarly, (**B**) shows the genome-wide expression on B-subgenome and (**C**) shows the expression on D-subgenome. The candidate genes have been shown inside the circos plots (**D**) Histogram representation of expression level of candidate genes and their homeologs w.r.t to two different temperature conditions: heat and cold stress. Brown and Light green color bars show the expression of genes w.r.t. 1 and 6h of heat treatment, pink and yellow color represent the expression of genes w.r.t 10 days and two weeks of cold treatment as provided by expVIP wheat genome expression browser. Log2 values were presented in the histograms.

Further, circos plots in Figure 6 present the genome-wide expression of all wheat genes on the chromosomes under heat stress for 1 and 6 hrs timepoints of two biological replicates (SRA accession no. GSE139019 with a window size of 5 MB). The location of candidate genes in the distal regions of chromosomes suggests the dynamic gene expression, as shown in Figure 6. In addition, we observed the dominant occurrence of H3K4me3 over the H3k9ac epigenetic marker in actively transcribing genes. Furthermore, H3K27me3, a repressive epigenetic marker, was found to be absent in a panel of 19 genes and their homeologs studied here (Figure 8). In a previous study (Wang *et al*. 2021), authors performed chromatin immunoprecipitation followed by sequencing (ChIP-seq) using H3K27me3 antibodies. However, they were unable to detect the presence of this repressive signature on downregulated wheat gene homeologs. Repressive markers H3K27me3, H3K9me3, H3K9me2, and H3K27me2 are also well-known. As a result, it is suggested that under specific stress situations, additional repressive marker(s) may be involved in downregulating or suppressing a specific homeolog of the bZIP gene instead of H3K27me3.

## Discussion

Because wheat is a low-cost cereal crop and a significant source of calories, understanding the regulatory mechanisms behind wheat’s capacity to withstand stress is essential. Since bZIP role in stress mitigation in plants is well known, this work focused on revealing and examining wheat bZIP TFs and their homeologs involved in temperature stress.

Cis-elements in the promoter are key to the gene regulation (Li *et al*. 2020). Promoter function analysis is performed by combining bioinformatics prediction and experimental validation. The type and number of multiple cis-elements in the promoter region of a gene can affect and may lead to divergent gene expression (Rizhsky *et al*. 2004). We analyzed the 2000 bp upstream region of candidate *bZIPs* using PlantCARE promoter prediction software. In the promoter region, in addition to some conserved elements, e.g. TATA and CAAT boxes, additional cis-elements were present, for example, Light-responsive elements G-Box, BOXI, ABA-responsive elements ABRE and MYC, ARE (anaerobic induction), GC-motif involved in anoxic specific inducibility, CGTCA-motif involved in the MeJa-response, G-box involved in light response and ABRE involved in the abscisic acid response, make a compelling reason for investigating their functional roles (Foster *et al*. 1994; Hirayama and Shinozaki 2010). These findings imply that TabZIP genes have a high propensity to respond to environmental stress and may control how plants respond to abiotic stresses. The density and diversity of various cis-elements suggest that activation of bZIPs under different biotic and abiotic stresses may depend on these respective DNA sequences.

Since bZIPs promoter regions are enriched with stress- and development-related cis-elements motifs (Ramírez-González *et al*. 2018) can be correlated to gene expression using publicly accessible data, such as RNA-seq tools (expVIP) (Borrill *et al*. 2016; Ramírez-González *et al*. 2018). In an earlier study, the bZIP genes were reported to be involved in temperature signalling and ABA response (Zhang *et al*. 2020). We observed that cis-acting elements such as LTR associated with cold temperature response, ARE, DRE, MYC, and MYBs were enriched in the promoters of TabZIPs (Figure 3).

It is observed that there is a significantly elevated gene expression levels in cold stress. Further, the overexpression protocol increased the resistance to cold stress conditions in transgenic plants (Shen *et al*. 1996; Finkelstein and Lynch 2000; Liang *et al*. 2022). TraesCS3D02G364900(ABI5), an attractive candidate gene for plant stress tolerance, exhibited a profound gene expression with a 9.22 LFC at 4 °C exposure. It was reported earlier that a significant gene expression level in cold stress and overexpression study increased the resistance to cold stress conditions in transgenic plants (Liang *et al*. 2022). It emerged as a candidate gene for the low-temperature response gene from the bZIP gene family in bread wheat. In this study novel, stress-related bZIPs TraesCS1A02G072600 showed 3.42 LFC at 4 °C stress. At 42 °C, it increased to 4.09 LFC and followed a similar expression trend of 5.62 LFC at 37 °C temperature. Like TraesCS1A02G072600 TraesCS6A02G096300, another novel TabZIP gene showed strong expression changes at different temperatures. At cold stress condition of 4 °C, it showed expression of 5.44 LFC. In the heat stress conditions, at 42 °C and 37 °C, the expression level increased by 9.41 LFC and 6.53 LFC, respectively. In conclusion, TabZIPs exhibit dynamic expression in response to various stress conditions.

Earlier studies in *Arabidopsis* demonstrated that in cold stress conditions, there is an enrichment in the H3K27me3 polycomb repressive complex 2, leading to the silencing of the Flowering locus C (FLC) expression (Xu *et al*. 2022). Wheat research has shown that under heat- and cold-stress conditions, localization of the chemically modified histone H2K4me3, H3K9ac, and H3K27me3 in promoter regions can result in either gene activation or repression (Ramírez-González *et al*. 2018; Wang *et al*. 2021). A closer examination of the physical co-ordinates of 19 temperature stress-related genes using the ChIP-seq data tracks for the presence of histone epigenetic markers H3K4me3, H3K9ac, and H3K27me3 revealed a mechanistic basis of asymmetric bZIP homeologs expression (Figure 8). The H3K4me3 occurrence is more than the H3K9ac. Although H3K9ac is expected to be populated in the promoter for the elevated gene expression, the H3K4me3 is required for engaging the TFIID for the acetylated histone to become functionally active (Clouaire *et al*. 2012).

In addition to TF-bound cis-elements and histone markers, chromosomal location influences gene activity. Distal genes are dynamic, and their position from a chromosome’s centromeric region determines its expression profile. It is observed that 19 temperature stress-related genes with their homeologs are all located at the distal end of the chromosomes, suggesting high dynamic activities of *TabZIP*s (Figure 6).

In the homeolog crossing-over event, there are specific amino acid changes in the sequence of most homeologs in hexaploid bread wheat. A single amino acid change may profoundly affect the nucleation of secondary structure formation in a bZIP. Figure 7 shows additional interactions when a homeolog has different amino acids. For example, A198100 has positively charged K at ‘c’ position of 4^th^ heptad that can interact with two negatively charged E at ‘g’ and ‘f’ positions (Simm *et al*. 2017). No such intramolecular interactions are expected in homeolog B117400, and D171700, where uncharged M is present instead of K. Such additional interactions may stabilise trigger sequences enabling the bZIP to preserve its structure in high or low temperatures (Figure 7). 5 bZIPs showed homeolog-specific amino acid changes that may lead to additional intramolecular interactions in a monomer. It will be interesting to perform biochemical and biophysical studies comparing different homeologs for their stability and specificity under temperature stress. Other 5 bZIPs showed amino acid change but appeared not to participate in additional interactions (Supplementary Figure 3). Since the protein sequences of 9 bZIPs are identical, epigenetic markers other than H3K4me3, H3K9ac, and H3K27me3 may be responsible for their varied expression pattern under stress (Figure 8). In conclusion, understanding asymmetric gene expression in wheat is important for developing new wheat varieties with improved agronomic traits and recognizing the fundamental biology of polyploid organisms. Based on the localization of *TabZIPs* on the distal regions of the chromosomes, this study, cis-element analysis, genetic and the epigenetic basis behind the asymmetric homeolog expression, helps to redeem the stress response by the specific bZIP candidates in *T aestivum*. The study may help to conduct further molecular and biochemical experiments to explore the capability of these novel and uncharacterized TabZIP proteins.

**Figure 7:**
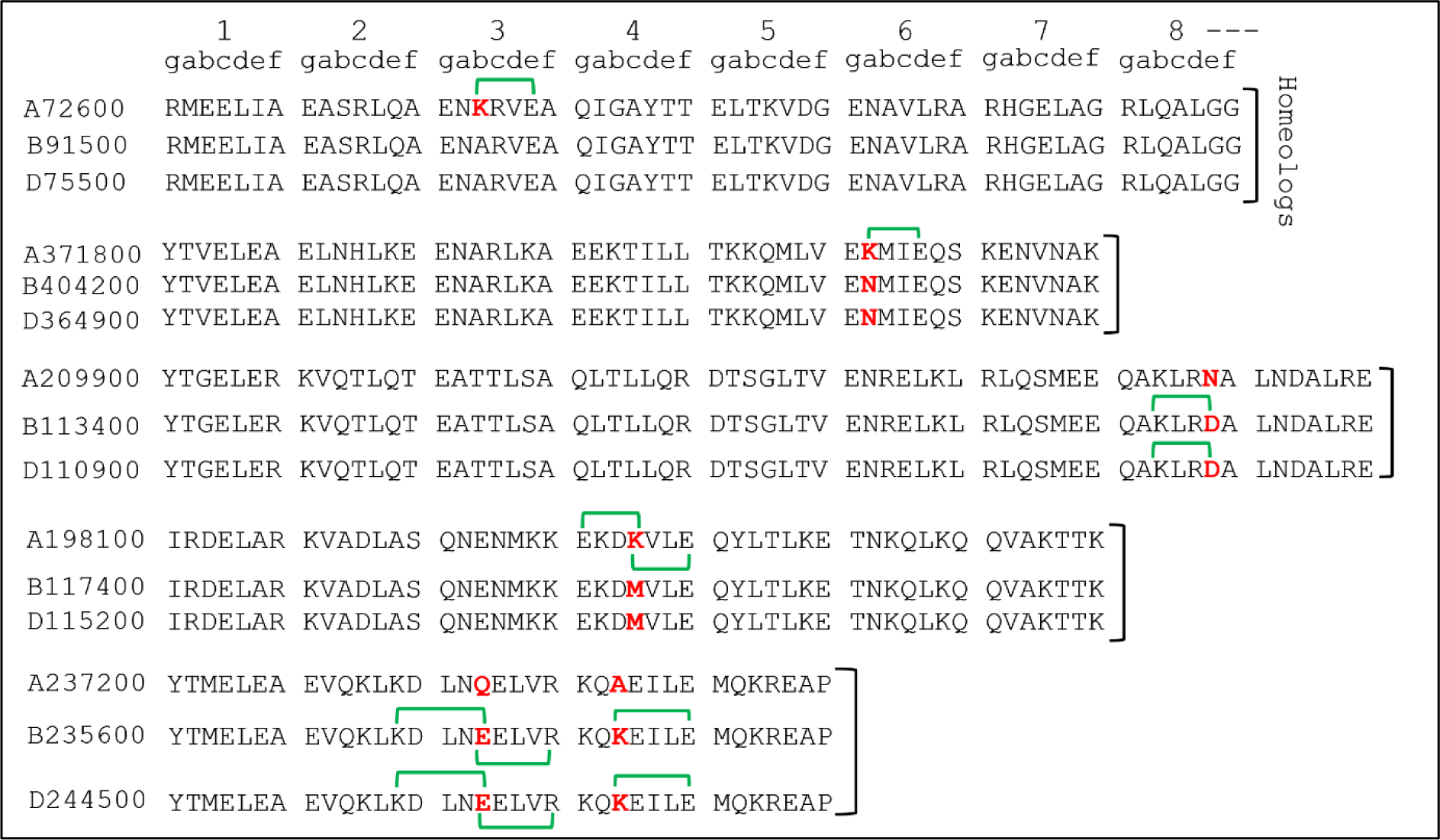
Amino acids sequences of three homeologs of 5 TabZIP proteins. The numbers at the left are the Unique identifiers in Treas IDs of stress-related wheat bZIPs. A, B, and D represent three subgenomes. Delineation of peptide sequences are shown as heptads (gabcdef) of seven amino acids. Numbers (1-8) shown at the top depict heptad length. In a few cases, for clarity, not all heptads are shown. Amino acids changes are shown in bold red letters. Additional interactions are shown as green brackets.

**Figure 8:**
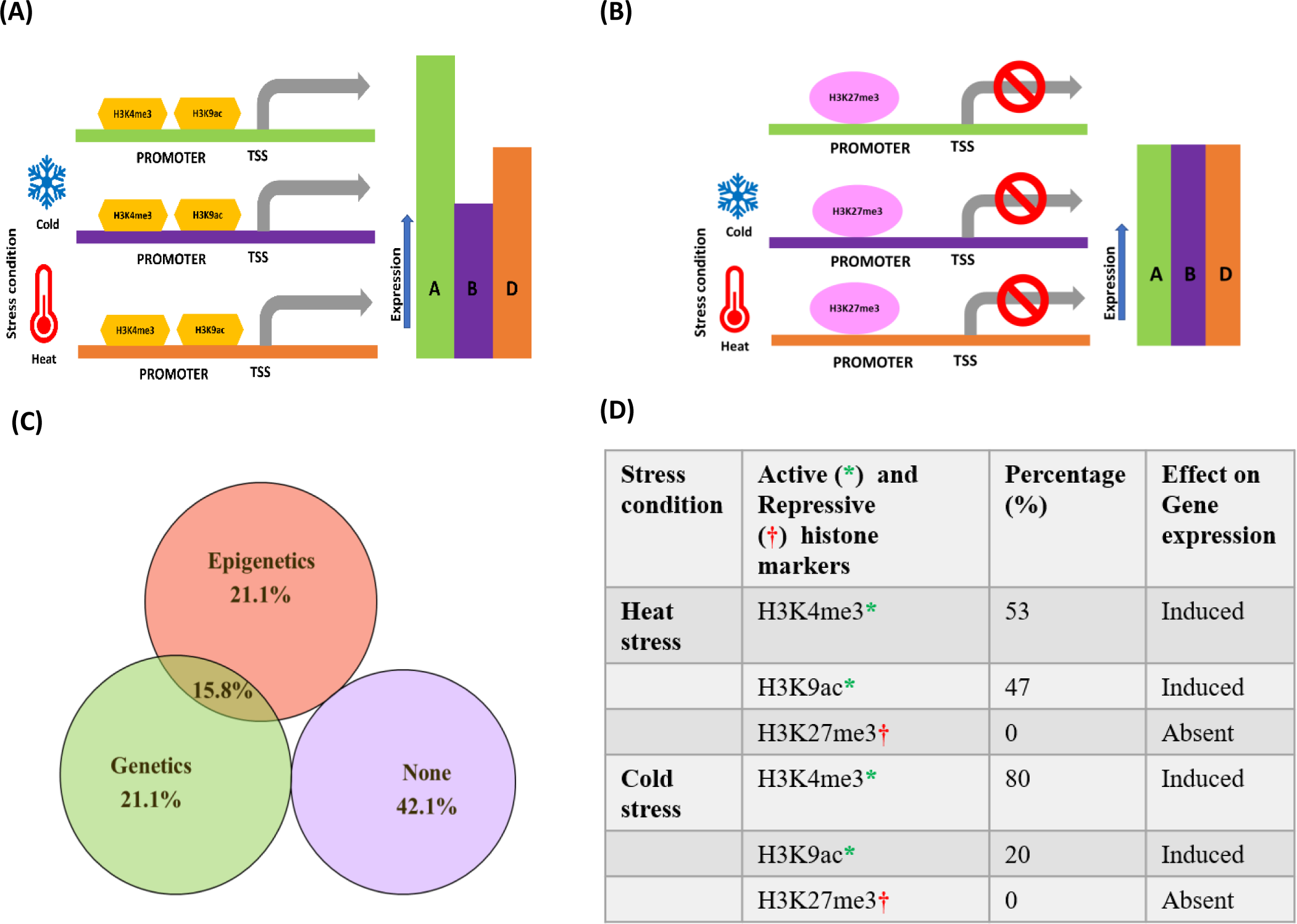
Abridged model reperesentation of epigenetic and genetic factors responsible for biased gene expression in *T.aestivum*. (**A)** The dominance and suppression in homeolog gene expression derived by the presence of active epigenetic histone markers such as H3K4me3, and H3K9ac leading to elevated and decreased gene expression in response to abiotic stress changes such as heat and cold temperature conditions. (**B)** The presence of repressive histone marker such H3K27me3 leads to non-significant and suppressed gene expression w.r.t temperature change. (**C)** Venn diagram showing the percentage of genetic, epigenetic (and overlapping) basis of asymmetric bZIP expression with 42.1 % unexplored coverage. (**D)** Table representing the details of epigenetic markers and effect on gene expression in the temperature stress conditions.

## Acknowledgments

We thank the Executive Director National Agri-Food Biotechnology Institute (NABI), Mohali, for research facilities and the Department of Biotechnology (DBT), New Delhi, for funding. RK acknowledges the Department of Science and Technology (DST) for the DST-Inspire fellowship.

## Competing Interest

Authors declare no competing interest.

## Conflict of Interest

Authors declare that there is no conflict of interest regarding publication of this research paper.

## Author contribution

RK and VR conceptualized the idea and designed the study. RK and VR designed the experiments, analysed the data and wrote the manuscript. RK conducted experiments with the help from SGD, HP, KK and AKP. RK prepared the figures and visualized the Bioinformatic data with the help from DS, PK and GK. All authors contributed in the editing of paper and have approved the final version of the manuscript.

## Notes

https://www.ncbi.nlm.nih.gov/geo/query/acc.cgi?acc=GSE139019

### Competing Interest Statement

The authors have declared no competing interest.

