## supplementary_figures for "Identifying the genetic and epigenetic basis for asymmetric bZIP expression in temperature-stressed bread wheat"

Website: www.nabi.res.in

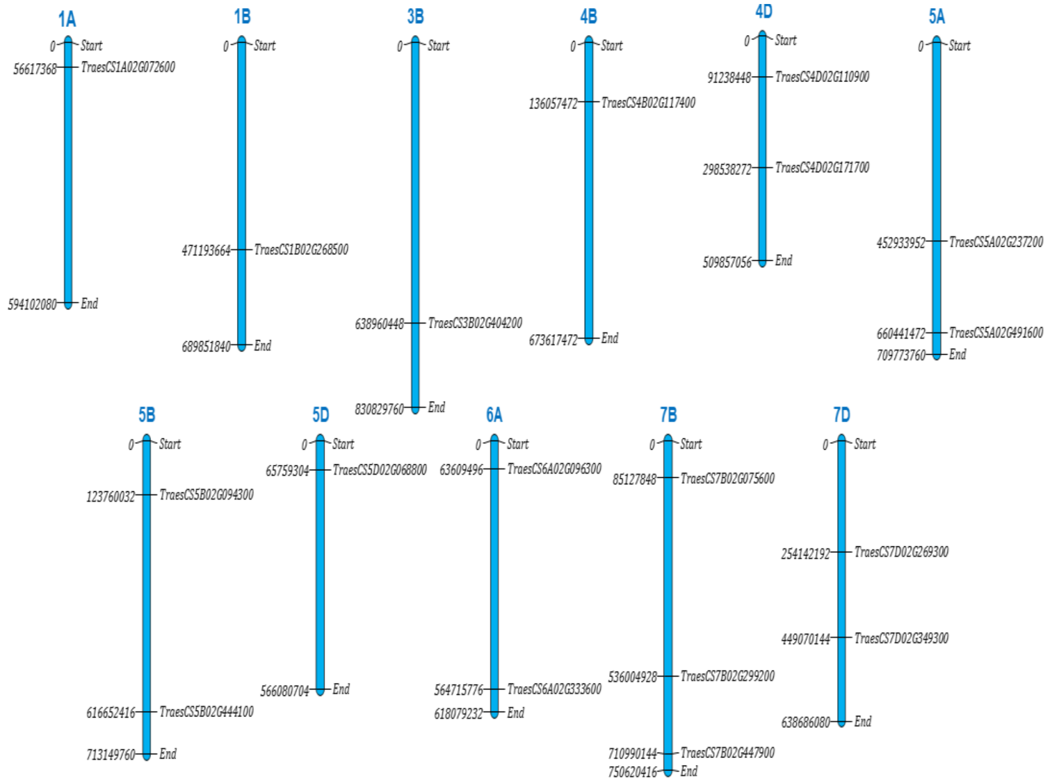


**Supplementary figure1**: Chromosome distribution of candidate bZIPs across A, B and D subgenomes of *Triticum aestivum*

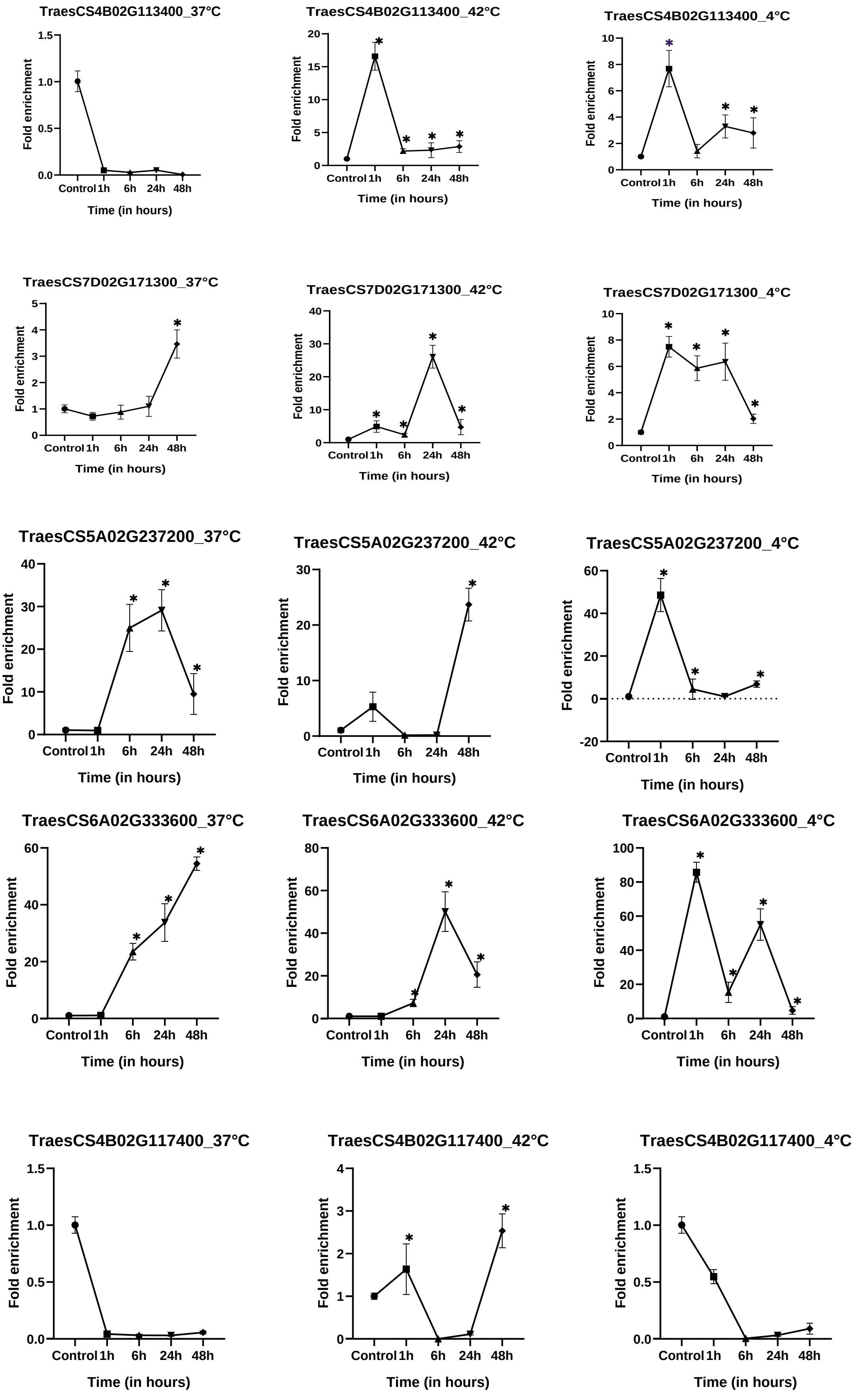

TraesCS4B02G113400_37°C
TraesCS4B02G113400_42°C
TraesCS4B02G113400_4°C
TraesCS7D02G171300_37°C
TraesCS7D02G171300_42°C
TraesCS7D02G171300_4°C
TraesCS5A02G237200_37°C
TraesCS5A02G237200_42°C
TraesCS5A02G237200_4°C
TraesCS6A02G333600_37°C
TraesCS6A02G333600_42°C
TraesCS6A02G333600_4°C
TraesCS4B02G117400_37°C
TraesCS4B02G117400_42°C
TraesCS4B02G117400_4°C
Fold enrichment
Control 1h 6h 24h 48h
Time (in hours)

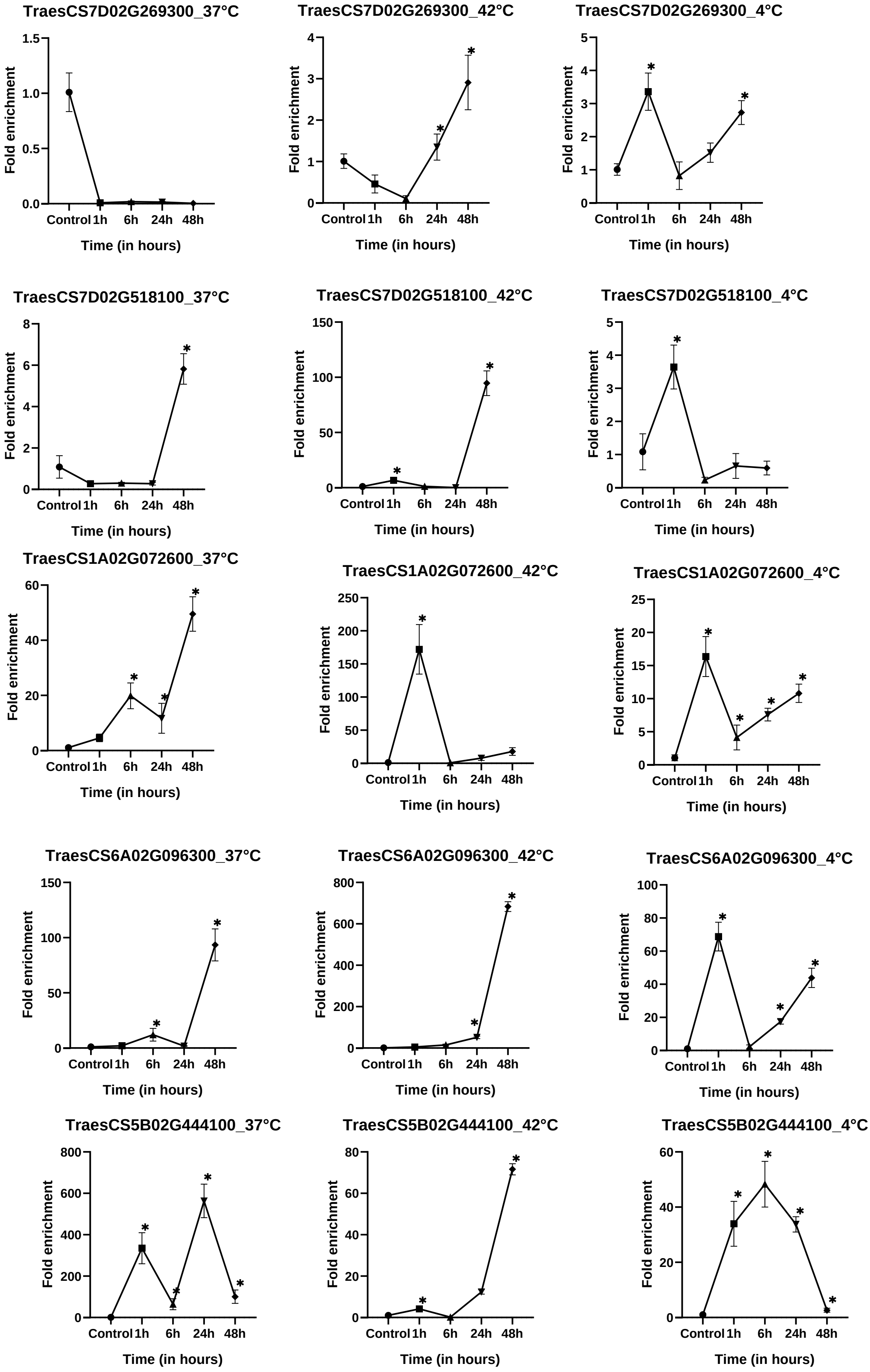
TraesCS7D02G269300_37°C
TraesCS7D02G269300_42°C
TraesCS7D02G269300_4°C
TraesCS7D02G518100_37°C
TraesCS7D02G518100_42°C
TraesCS7D02G518100_4°C
TraesCS1A02G072600_37°C
TraesCS1A02G072600_42°C
TraesCS1A02G072600_4°C
TraesCS6A02G096300_37°C
TraesCS6A02G096300_42°C
TraesCS6A02G096300_4°C
TraesCS5B02G444100_37°C
TraesCS5B02G444100_42°C
TraesCS5B02G444100_4°C
Fold enrichment
Time (in hours)
Control
1h
6h
24h
48h
*

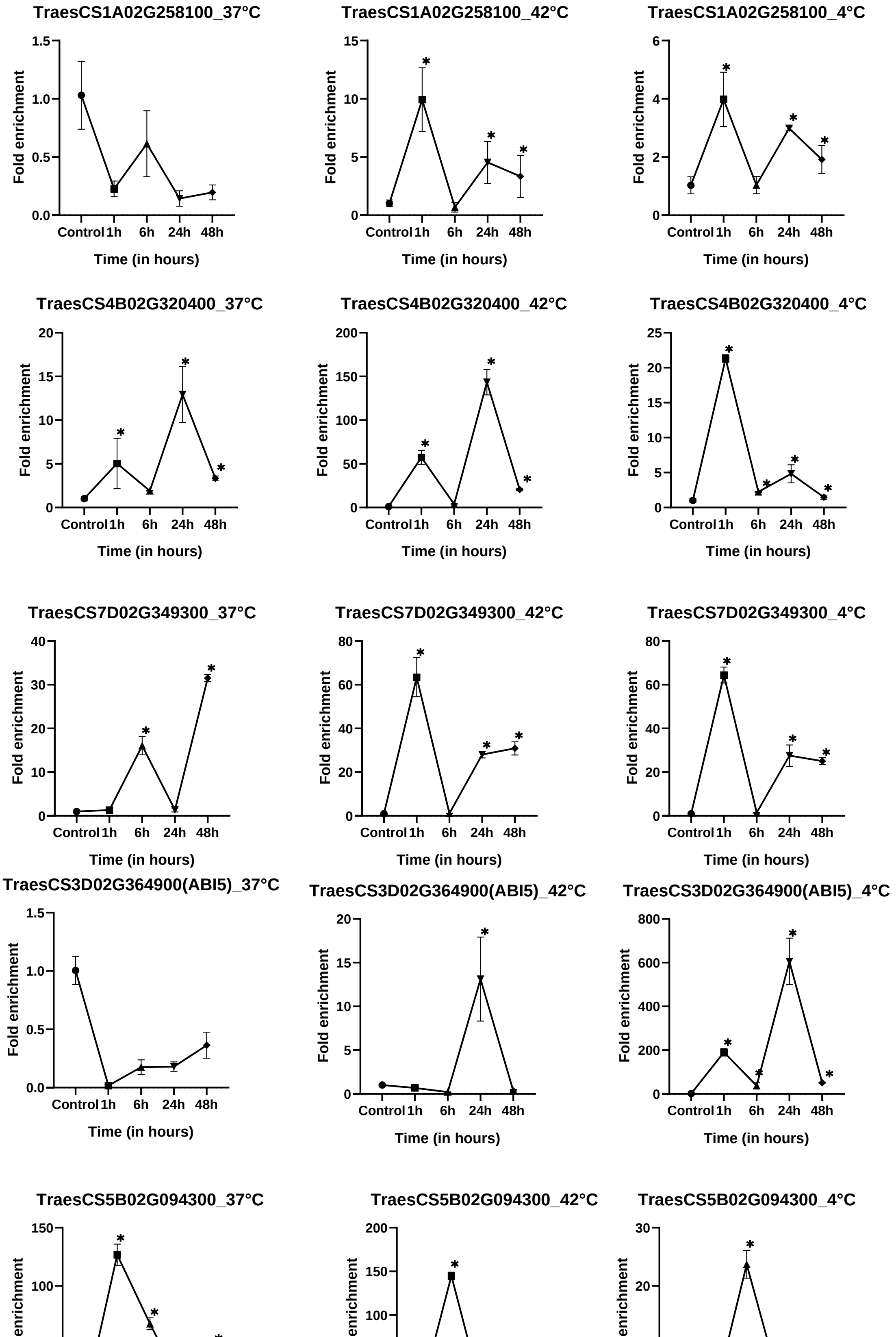
TraesCS1A02G258100_37°C
TraesCS1A02G258100_42°C
TraesCS1A02G258100_4°C
TraesCS4B02G320400_37°C
TraesCS4B02G320400_42°C
TraesCS4B02G320400_4°C
TraesCS7D02G349300_37°C
TraesCS7D02G349300_42°C
TraesCS7D02G349300_4°C
TraesCS3D02G364900(ABI5)_37°C
TraesCS3D02G364900(ABI5)_42°C
TraesCS3D02G364900(ABI5)_4°C
TraesCS5B02G094300_37°C
TraesCS5B02G094300_42°C
TraesCS5B02G094300_4°C
Fold enrichment
Control
1h
6h
24h
48h
Time (in hours)
*

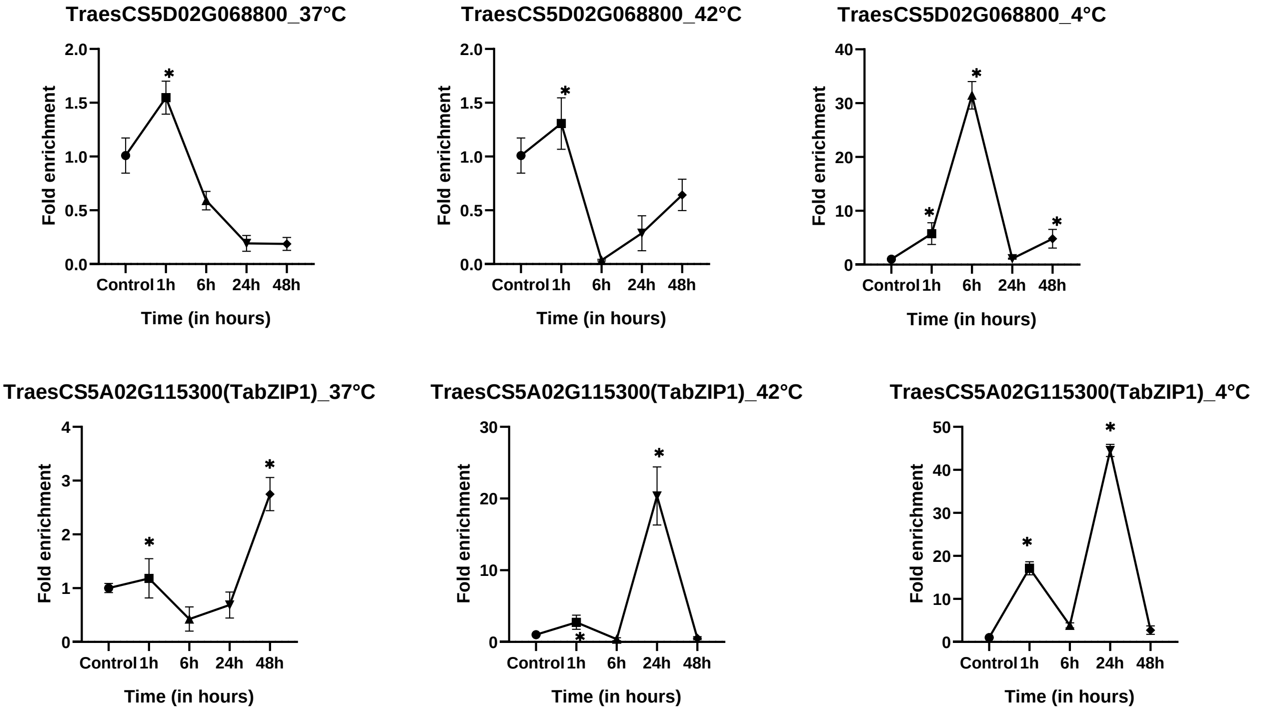


**Supplementary Figure 2:** Quantitative real time PCR-based expression validation of candidate genes at three temperature stress conditions 37 ˚ C and 42 ˚ C and 4˚ C, (1 gene in each lane) with 1,6,24 and 48 h time-points. Student's t-test was employed for plotting the graphs by GraphPad-Prism. Significance level was represented by * depicting the *p*-value < 0.05

| | 1 | 2 | 3 | 4 | 5 | 6 | 7 | 8 | 9 |
|---|---|---|---|---|---|---|---|---|---|
| | gabcdef | gabcdef | gabcdef | gabcdef | gabcdef | gabcdef | gabcdef | gabcdef | gabcdef |
| A268700 | YIQELEH | KVQVLQT | EATTLSA | QLTMMQR | | | | | |
| D269300 | YIQELEH | KVQVLQT | EATTLSA | QLTMMQR | | | | | |
| B075600 | YIMELEA | EVAKLKE | NNEALQK | KQVEMLQ | KQKDEVI | ERIEKQL | | | |
| A170600 | YIMELEA | EVAKLKE | NNEALQK | KQVEMLQ | KQKDEVI | ERIEKQL | | | |
| D171300 | YIMELEA | EVAKLKE | NNEALQK | KQVEMLQ | KQKDEVI | ERIEKQL | | | |
| A530300 | HTALLEE | EVAHLKA | VNQQLVK | KLQSHSA | LEAEVAR | LRCLLVD | IRGRIEG | | |
| B447900 | HTALLEE | EVAHLKA | VNQQLVK | KLQSHSA | LEAEVAR | LRCLLVD | IRGRIEG | | |
| D518100 | HTALLEE | EVAHLKA | VNQQLVK | KLQSHSA | LEAEVAR | LRCLLVD | IRGRIEG | | |
| A398400 | YVKDLET | **R**SKYLEA | ECRRLSY | ALQCCAA | ENMALRQ | NMLKDRP | | | |
| B299200 | YVKDLET | **K**SKYLEA | ECRRLSY | ALQCCAA | ENMALRQ | NMLKDRP | | | |
| D392800 | YVKDLET | **K**SKYLEA | ECRRLSY | ALQCCAA | ENMALRQ | NMLKDRP | | | |
| A258100 | YVEELEE | KVKSMNS | VINDLNS | KISFIVA | ENATLRQ | QLGSGG | | | |
| B268500 | YVEELEE | KVKSMNS | VINDLNS | KISFIVA | ENATLRQ | QLGSGG | | | |
| D257400 | YVEELEE | KVKSMNS | VINDLNS | KISFIVA | ENATLRQ | QLGSGG | | | |
| A491600 | YIVELEQ | KVQILQT | EATTLAA | QINLLQR | DSSAVAT | QNNELRF | RLQAMEQ | QAQLRDA | LN**E**ALTG |
| B320400 | YIVELEQ | KVQILQT | EATTLAA | QINLLQR | DSSAVAT | QNNELRF | RLQAMEQ | QAQLRDA | LN**E**ALTG |
| D316900 | YIVELEQ | KVQILQT | EATTLAA | QINLLQR | DSSAVAT | QNNELRF | RLQAMEQ | QAQLRDA | LN**D**ALTG |
| A135600 | YMTELER | KVQTLQT | EATTLSA | QLTLFQR | DTTGLSS | ENAELKI | RLQAMEQ | QAQLRDA | LNDALKQ |
| B169600 | YMTELER | KVQTLQT | EATTLSA | QLTLFQR | DTTGLSS | ENAELKI | RLQAMEQ | QAQLRDA | LNDALKQ |
| D171700 | YMTELER | KVQTLQT | EATTLSA | QLTLFQR | DTTGLSS | ENAELKI | RLQAMEQ | QAQLRDA | LNDALKQ |
| A440400 | HLNELEA | QVSQLRV | ENSSLLR | RLADVNQ | KYNGAAV | DNRVLKA | DVETLRA | KVKMAED | SVKRVTG |
| B444100 | HLNELEA | QVSQLRV | ENSSLLR | RLADVNQ | KYNGAAV | DNRVLKA | DVETLRA | KVKMAED | SVKRVTG |
| D447500 | HLNELEA | QVSQLRV | ENSSLLR | RLADVNQ | KYNGAAV | DNRVLKA | DVETLRA | KVKMAED | SVKRVTG |
| A088300 | HLDELVQ | EVARLKA | ENARVLA | RANDITG | QFVRVDQ | ENTVLRA | RAAELGD | RLRSVNQ | VLRVVEE |
| B094300 | HLDELVQ | EVARLKA | ENARVLA | RANDITG | QFVRVDQ | ENTVLRA | RAAELGD | RLRSVNQ | VLRVVEE |
| D100700 | HLDELVQ | EVARLKA | ENARVLA | RANDITG | QFVRVDQ | ENTVLRA | RAAELGD | RLRSVNQ | VLRVVEE |
| A057500 | HQNDIES | QVTQLRA | **E**NASLLK | RLTDMTQ | KYKEA**S**L | GNRNLTV | DIETMRR | KVNIAEE | AVRRVTG |
| B059200 | HQNDIES | QVTQLRA | **D**NASLLK | RLTDMTQ | KYKEA**F**L | GNRNLTV | DIETMRR | KVNIAEE | AVRRVTG |
| D068800 | HQNDIES | QVTQLRA | **E**NASLLK | RLTDMTQ | KYKEA**S**L | GNRNLTV | DIETMRR | KVNIAEE | AVRRVTG |
| A115300 | ECEELAQ | RAEVLKQ | ENASLKD | EVSRIRK | EYDELLS | KNSSLKD | NVGDKQH | KTDEAGL | DNKLQHS |
| B122000 | ECEELAQ | RAEVLKQ | ENASLKD | EVSRIRK | EYDELLS | KNSSLKD | NVGDKQH | KTDEAGL | DNKLQHS |
| D124600 | ECEELAQ | RAEVLKQ | ENASLKD | EVSRIRK | EYDELLS | KNSSLKD | NVGDKQH | KTDEAGL | DNKLQHS |
| A333600 | YMMELET | EVAKLKE | RNEELQR | KQAE**M**LE | RQKNEVF | EKVTRQA | | | |
| B364000 | YMMELET | EVAKLKE | RNEELQR | KQAE**I**LE | RQKNEVF | EKVTRQA | | | |
| D312800 | YMMELET | EVAKLKE | RNEELQR | KQAE**I**LE | RQKNEVF | EKVTRQA | | | |
| A096300 | HLDDLAA | QAAHLRR | ENAHVAA | ALGLTAR | GLQAVDA | ENAVLRT | QAAELAA | RLHSLND | IIACMSA |
| B124700 | HLDDLAA | QAAHLRR | ENAHVAA | ALGLTAR | GLQAVDA | ENAVLRT | QAAELAA | RLHSLND | IIACMSA |
| D087400 | HLDDLAA | QAAHLRR | ENAHVAA | ALGLTAR | GLQAVDA | ENAVLRT | QAAELAA | RLHSLND | IIACMSA |
| A373800 | YMGELE**A** | KVKDLET | RNSELEE | RLSTLQN | ENQML**R**Q | ILKNTTG | | | |
| D349300 | YMGELE**V** | KVKDLET | RNSELEE | RLSTLQN | ENQML**K**Q | ILKNTTG | | | |

**Supplementary Figure 3:** Amino acids sequences of three homeologs of 14 bZIP proteins. The numbers at the left are the Unique identifiers in Traes IDs of stress-related wheat bZIPs. A, B, and D represent three subgenomes. Delineation of peptide Sequences are shown as heptads (gabcdef) of seven amino acids. Numbers (1-8) shown at the top depict heptad length. In a few cases, for clarity, not all heptads are shown. Amino acids changes are shown in bold red letters.

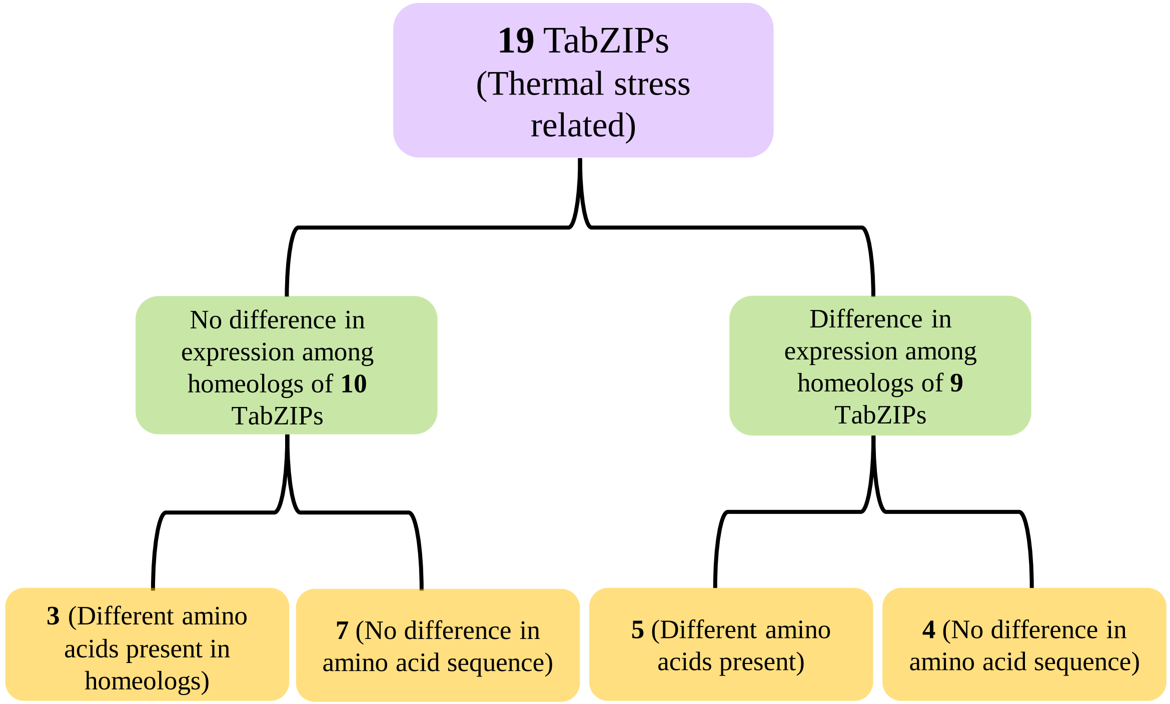


**Supplementary Figure 4:** Depiction of 19 temperature- stress related TabZIPs into two sections with change and no-change in temperature induced expression pattern in homeologs. Further, these two categories contained 3 and 5, 7 and 4 number of TabZIPs where amino acid differences were observed or remain unchanged in the sequence.

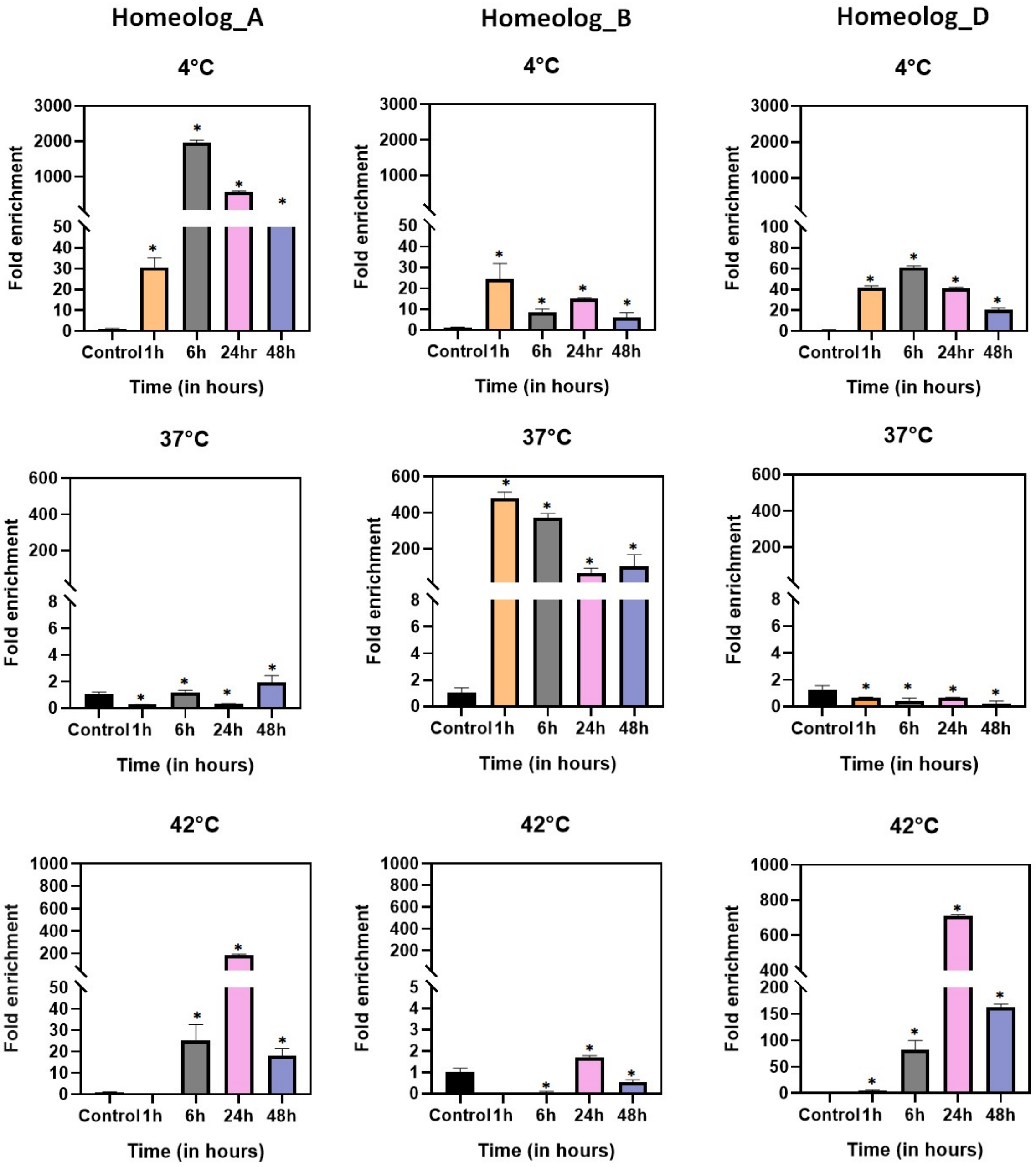


**Supplementary Figure 5:** Quantitative real time PCR-based asymmetric homeolog expression validation between three homeologs (TraesCS1A02G072600, TraesCS1B02G091500, and TraesCS1D02G075500) at three temperature stress conditions 37˚C and 42˚C and 4˚C, (1 temperature condition in each lane) with 1,6,24 and 48 h time-points. Student's t-test was employed for plotting the graphs by GraphPad-Prism. Significance level was represented by * depicting the $p$-value < 0.05
